# Atomic-level characterization of conformational transition and substrate binding of xCT transporter

**DOI:** 10.1101/389643

**Authors:** M. Sharma, A. C. Rohithaswa

## Abstract

xCT is a component of heterodimeric amino acids transporter system Xc- that has been known to work at the cross-roads of maintaining neurological processes and regulating antioxidant defense. xCT is a sodium-independent amino acid antiporter, that imports L-cystine and exports L-glutamate in a 1:1 ratio. The transporter has 12 transmembrane domains with intracellular N- and C-termini, which can undergo various conformational changes while switching the ligand accessibilities from intracellular to extracellular site. In the present study, we generated two homology models of human xCT in two distinct conformations: inward facing occluded state and outward facing open state. We investigated the conformational transitions within these two states by employing series of targeted molecular dynamics simulations. Our results indicated the substrate translocation channel composed of transmembrane helices TMs 1, 3, 6, 8, and 10. Further, we analyzed the ligand binding within the intermediate conformations obtained from the transition simulations. We docked anionic L-cystine and L-glutamate within the cavities alone or in combination to assess the two distinct binding scenarios for xCT as antiporter. We also assessed the interactions between the ligand and xCT and observed that ligands bind to similar residues within the channel, and these residues are essential for substrate binding/permeation. In addition, we analyzed the correlations between ligand binding and conformational transition and observed conformations that are representatives for intermediate ligand bound states. The results presented in the study provide insights into the interplay of conformational transition and ligand binding as xCT goes from one probable conformation to another while transporting the ligand. And the data thus adds to the existing evidence of alternating access mechanism pertaining to the functioning of transporters.

## Introduction

The cystine-glutamate transporter (xCT) is a component of heteromeric, sodium-independent amino acid transport (HAT) system, Xc- with high specificity for cystine and glutamate (1, 2). The function of Xc- primarily connects neurotransmission and behavior with the antioxidant defense(3–5). System Xc- includes specific light chain transporter xCT (~40kDa) and heavy chain, 4F2hc (~80kDa) which are linked by a disulfide bridge(6, 7). In a functional dimer, 4F2hc is responsible for trafficking of the light chain and xCT is required for the transport activity of the dimer. Reconstitution studies on similar HAT system (SLC7A9) showed that light chains are fully functional in the absence of its corresponding heavy subunit(8). Expression of xCT on the cell membrane is essential for the uptake of cystine required for intracellular glutathione (GSH) synthesis and maintaining the intracellular redox balance(9). Extracelullar glutamate acts as a competitive inhibitor for cystine uptake via system Xc-, indicating that xCT primarily imports anionic cystine and exports glutamate in 1:1 ratio as an electroneutral antiporter exchange system for cystine/glutamate(10). Impairment of xCT results in disruption in glutamate homeostasis as observed in primary gliomas, resulting in elevated glutamate secretion and neuronal cell death(11–13). Dysfunction of xCT or decreased levels of mRNA expression of SLC7A11, gene associated with encoding xCT has been linked with various neurodegenerative disorders(5, 14, 15), such as schizophrenia(16), amyotrophic lateral sclerosis (ALS)(17), multiple sclerosis (MS)(18), and Parkinson’s diseases(19). On the other hand, xCT is upregulated in various cancer cell lines under oxidative stress including pancreatic cancer(20), breast cancer (21, 22), bladder carcinoma cells (23), and lung tumor progression(24). Expression of xCT has also been proposed as a predictor of disease recurrence in patients with colorectal cancer(25). Enhanced biosynthesis of intracellular GSH via xCT protects cancer cells from drug-induced oxidative stress by mediating drug detoxification or inactivation(26–28). In addition, xCT has been identified as the predominant mediator of Kaposi’s sarcoma-associated herpesvirus (KSHV) fusion and entry permissiveness into cells(29–31). These findings make xCT a promising therapeutic target in cancer therapy(32–35) and in neurodegenerative or neuroinflammatory diseases(3, 5, 36, 37).

A detailed molecular understanding of conformational transitions of xCT during the substrate transport would offer new opportunities for drug discovery(38, 39). This warrants the need to have structural information of human cystine transporter. There have been structural experimental studies (40–42) indicating that xCT is a protein with 12 transmembrane (TM) helices, both N- and C- termini located intracellularly, and re-entrant loop between helices TM2 and TM3 that participate in substrate permeation. The only structures available till date are two homology models. One model was constructed using ApcT crystal structure(14). Another xCT modeled structure was reported recently (43) where human xCT structure was mapped onto the modeled structure of homologous fungal cystine transporter, CgCYN. With the advancement in membrane structural biophysics and crystallography, the transporters have been captured in different conformational states, suggesting that there are several conformational states accessible to these transporters as they switch accessibility from one side of the membrane to the other (44–52), and are now thought to operate by a common “alternating access” model(53, 54). This implies that there are still fundamental questions that are open to investigate further about xCT. What are the other conformations possible for xCT based on the structural data available? Are there intermediate conformations between these states providing insights about the conformational transitions of xCT or the substrate movement across the membrane? In this study, we have constructed homology model of xCT in two distinct conformations: inward facing occluded state and outward facing open state. More detailed investigations of structural rearrangements of transporters at atomic level have been made possible by molecular dynamics simulations (55–61). However, these conventional simulations lack to capture critical events pertaining either to conformational transitions between two conformations, or to substrate permeation pathway(60, 62) since such events require longer timescales. Fortunately, the timescale problem faced by atomic-level simulations can be alleviated by relying on biased simulations where artificial forces are generally used to accelerate transitions(63–68). One such technique, targeted molecular dynamics (TMD) simulations studies have previously been carried out to understand the conformational transitions of ATP-binding cassette transporter, BtuCD(69), human Glucose transporter GLUT1(70), Potassium channels (71), AcrB Efflux transporter (72). In this study, we have modeled xCT is two distinct conformations and employed series of TMD simulations to understand the conformational transition pathway between these two conformations. The intermediates were identified, and probable ligand binding sites were explored with both anionic L-cystine and L-glutamate substrates. The substrate bound studies provided insights about the substrate translocation/permeation pathway.

## Methods

### Identification and modeling of transmembrane (TM) region of xCT

Human xCT sequence was retrieved from NCBI, and an initial topology prediction was carried out using Constrained Consensus TOPology (CCTOP) server(73), which utilizes ten topology prediction methods: HMMTOP, MemBrain, Memsat, Octopus, Philius, Phobius, Pro, Prodiv, Scampi, ScampiMsa, and TMHMM in combination with the topology information from PDBTM, TOPDB, and TOPDOM databases using the probabilistic framework of hidden Markov model (Fig SI1). xCT was predicted to be a 12-transmembrane helical protein with N- and C- termini located inside the cell. Sequence spanning transmembrane region (residues 45 to 512) was submitted to HHPred server against PDB structures for homology detection, and two templates, ApcT (PDb id:3GIA)(74) and AdiC (PDb id:5J4I)(75) were selected using HHblits multiple sequence alignment method(76). Transmembrane region of xCT showed ~51% sequence similarity with both ApcT (E value:2.2e-32) and AdiC (E value:1.9e-32) (Fig S12). These two structures are in different conformational states of their transport cycle. AdiC arginine antiporter is reported as substrate free outward facing state, and ApcT is reported as substrate-free inward facing occluded open state. The only inward open conformation known for any amino acid transporter is that of GadC(77). However, the reported structure consists of C-terminal fragment (C-plug) whose displacement is requisite for GadC transport activity. In absence of such information for xCT, we decided not to opt for GadC as template for this study. Comparative modeling of TMD region of xCT spanning residues 45 to 512 including TMD helices and connecting loops was carried out based on multiple sequence alignment of these two templates using Modeller9v15(78). Two representative models corresponding to the two templates were selected from the collection of 5000 models built by Modeller, based on the evaluation of discrete optimized protein energy (DOPE) potential. The models were further subjected to loop refinement module of Modeller9v15 to refine the connecting intracellular and extracellular loops, and energetically favorable models were selected among the 1000 generated models based on the DOPE score. Thus, two models of xCT have been generated using two different templates in different conformational states of access cycle herein referred as Model-Cioc (template structure: 3GIA), and Model-Cout (template structure: 5J4I) (Fig. S13).

### Minimization and refinement of modeled structure using MD in explicit membrane

The selected models were minimized further to obtain stable structures using Molecular dynamics simulations. Simulation setup with model inserted into a heterogeneous fully hydrated bilayer of size 85Åx85Å, consisting of lipids as well as cholesterol molecules, was obtained using CHARMM-GUI(79) using CHARMM36m forcefields(80) for lipids as well as protein. Prior to membrane insertion, we calculated rotational and translational positions of xCT models in membranes using PPM server(81), and the obtained orientation was used for positioning within lipid bilayer in CHARMM-GUI. The constituents of lipid bilayer were selected in order to mimic the human brain barrier membrane environment(82) - DPPC(5%):POPC(20%):POPE(15%):POPS(5%):POPI(5%):PSM(30%) lipids with cholesterol (20%). The structures were then solvated with TIP3P water molecules, followed by addition of K+ and Cl^-^ ions for 0.15M concentration. The assembled simulation system consisted of ~90,000 atoms. The biomolecular systems were simulated using NAMD (83). Minimization was carried out for 10000 steps, and system was equilibrated for 0.5 ns, while slowly releasing the collective variable restraints to facilitate stable simulation. Finally, structure was further simulated without any restraints for 5ns at a constant temperature of 303K using Langevin thermostat with damping coefficient of 1 ps^−1^, and constant pressure of 1atm using semi-isotropic Nose-Hoover Langevin piston pressure control with oscillation period of 0. 05ps and oscillation decay time of 0.025ps. The van der Waals interactions were smoothly switched off at 10–12Å by a force-switching function, and long-range electrostatic interactions were calculated using particle mesh Ewald method(84). All bond lengths involving hydrogen atoms were fixed using SHAKE algorithm, enabling the use of 2fs time step. The final obtained structures were then used for targeted molecular dynamics.

### Targeted molecular dynamics between two modelled conformations

Targeted Molecular Dynamics (TMD) simulation(85) was employed to accelerate the conformational transitions by means of steering forces. The form of the force applied is 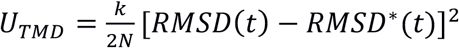 where *RMSD(t)* is the root mean square deviation of N atoms with respect to target structure at time *t*; and *RMSD*(t)* is the prescribed root mean square deviation value at time *t*. Series of targeted molecular dynamics (TMD) simulations were carried out using NAMDv2.12 to view the transition dynamics of xCT transporter from one conformational state to another. Models in inward occluded state and outward open states were considered as endpoints for both cycles of TMD. One cycle of TMD comprises of two sets of independent simulations: 10 short simulations with transition occurring within 25ns, and 3 longer simulations as independent runs with transition occurring within 200ns. For shorter and longer run simulations, force constants of 5 kcal.mol^−1^Å^−2^ and 2 kcal.mol^−1^Å^−2^, respectively per Ca atom for residues 37 to 473 was used to force them into the target structure with final rmsd as 0Å. Structure obtained at the end of each TMD run was further simulated for 5 ns. Similar protocol as used in above conventional MD was adopted for carrying out all the TMD simulations. In addition, harmonic restraints with force constant 10 kcal/mol were applied to restraint the orientation of overall protein molecule using *colvars* (collective variables) module. Table 2 summarizes the set of simulations carried out in this study. The total targeted molecular simulation time for xCT amounts for 1.7 μs.

**Table 2.** Targeted molecular simulations carried out in study

| Transition Cycle | TMD | Set I |  | Set II |  |
| --- | --- | --- | --- | --- | --- |
|  |  | Runs | Time | Runs | Time |
| Cycle I | Model_Cioc → Model_Cout | 10 | 25ns | 3 | 200ns |
| Cycle II | Model_Cout → Model_Cioc | 10 | 25ns | 3 | 200ns |

**Table 2.** Ligand docked xCT simulated systems

| S.No. | Ligand Docked | Docked structures | Time per run(ns) | Total simulation (ns) |
| --- | --- | --- | --- | --- |
| 1 | L-cystine | 45 | 20 | 900 |
| 2 | L-glutamate | 52 | 20 | 1040 |
| 3 | L-cystine +<br>L-glutamate | 15 | 25 | 375 |

### Generation of transition pathway and obtaining representative structures

In order to observe the underlying dynamics of TMD, we carried out principal component analyses using Gromacs2016(86) to observe the conformation sampling of all the trajectories during the two cycles. We pooled in all the conformations for both cycles by concatenating the trajectories, and calculated the eigenvectors using *covar* module, and projected the conformations of each cycle onto these eigenvectors using *anaeig* module. All heavy atom protein coordinates from the TMD trajectories obtained during both the cycles were submitted for clustering analysis using kmeans algorithm of *cpptraj* (87) module of AMBERTools17(88). Clustering was carried out on C□ atoms of residues 45 to 472 till 10 clusters are obtained and representative structures best fit to the cluster averaged structure were also obtained. These representative structures were used for docking purposes later.

### Docking of modeled structures with ligands

Coordinates of 10 representative structures obtained after clustering were used as receptor structures for docking. For ligands, coordinates for anionic L-cystine and L-glutamate were obtained from PubChem database (https://pubchem.ncbi.nlm.nih.gov) in sdf format. Binding site analysis was done by docking using AutodockVina(89), a computational docking program installed as plugin in PyMol(90). Since xCT is an antiporter, simultaneous efflux of glutamate and influx of anionic cystine is also possible. Therefore, we considered docking substrates first as single ligand docking and then docked both ligands.

#### (a) Docking with either L-cystine or L-glutamate

Prior to docking, cavities within the representative structures were analyzed using HOLE program(91). The docking box sizes for each model were selected such that the cavity within the central channel was covered within the box volumes of ~30Åx30Åx30Å enclosing the binding pocket. The residues within the binding pocket were defined as flexible residues to accommodate the ligands. 50 docked poses were generated per docking calculation and unique docked poses based on autodock vina score and manual inspection were filtered out for further simulations. For anionic cystine, there were 45 unique binding poses observed and for glutamate there were 52 binding poses observed. These single-ligand docked xCT conformations were then embedded in lipid bilayer as discussed above and simulated using similar protocol as for modeled structures for 20ns. Parameters for anionic cystine and glutamate were obtained for ligand generator module (92) of Charmm-GUI.

#### (b) Docking of both L-cystine and L-glutamate

The unique structures docked with single ligand (either anionic cystine or glutamate) obtained above were further used for blind docking with another ligand using Autodock Vina. For instance, 45 anionic cystine bound xCT conformations were docked again with glutamate ligand. Similarly, 52 glutamate bound xCT conformations were docked again with anionic cystine. For dual ligand docking, the box sizes were selected such that the cavities analyzed using HOLE program were covered within box volumes of ~30Åx30Åx30Å enclosing the binding pocket. Again 50 poses with both anionic cystine and glutamate docked ligands were generated for both the docking calculations. All these docked structures were then clustered using density-based clustering algorithm, *dbscan* implemented in *cpptraj* module of AmberTools17, with minimum cluster points as 20 and distance cutoff of 0.9Å. As distance metric, rmsd was calculated for Ca atoms of residues 45 to 472 and heavy atoms belonging to anionic cystine and glutamate. There were 15 unique docked conformations observed consisting both anionic cystine and glutamate. These xCT conformations were embedded in lipid bilayer and simulated for 20ns using similar protocol as for modelled xCT conformations. The ligand docked xCT simulated systems are summarized in Table 2.

### Interactions of ligands with protein

All the ligand docked conformations sampled at every 2ps were pooled in to visualize protein-ligand interactions in single ligand and dual ligand docked conformations. Interaction of ligand with protein residue was defined for when heavy atom of ligand interacts with protein residue within cutoff distance of 4.0Å. Variations in interactions between ligands and protein residues were analyzed using VMD1.9.3(93).

### Generation of substrate translocation pathway

For defining the substrate translocation pathway, all simulated conformations of the ligand docked structures were pooled in for translocation analysis. In order to characterize the relation between ligand placement along the channel and xCT conformation during the transition pathway, we defined two parameters: Z-direction of distance of ligand from the center of channel; and normalized root mean square distance of xCT from the two end points. For the former parameter, center of the translocation channel was defined by the center of cavity formed by TM helices 1, 3, 6, 8, and 10. The distance of center of mass of ligand to the center of translocation channel along the z direction was analyzed to define the position of ligand along the transporter. Since all the xCT conformations were first aligned to origin, the negative z- values will mean ligands placed towards the intracellular site and positive z- values will mean ligands placed towards the extracellular site. For the dual ligand docked conformations, this parameter relates to the z-component of distance between center of masses of the two ligands. For the second parameter, we calculated root mean square deviations of the translocation channel defined by helices TM 1,3,6,8, and 10 with respect to Model-Cioc and Model-Cout. Then the former rmsd value was subtracted from the latter and normalized over range −1:+1 for all the ligand docked conformations. This provided qualitative picture such that conformations in the negative range will have translocation channel similar to Model-Cioc, and conformations with normalized positive difference will have translocation channel similar to Model-Cout. All the conformations were clustered based on these two parameters using density based (dbscan) clustering program (94)implemented in R. DBSCAN is a partitioning method, and can find out clusters of different shapes and sizes from data containing noise and outliers. The algorithm requires to specify the optimal eps value and minimum points parameter, MinPts. The optimal eps parameter required for each clustering was determined by computing the k-nearest neighbor distance in a matrix of points. The respective conformations best fit to the centroids were also obtained.

### Calculation of water molecules within the substrate translocation pathway

We computed total number of water molecules present within the translocation pathway in presence of ligands using VMD1.9.3. The variation of number of water molecules with respect to the normalized rmsd difference as defined above was analyzed. To ensure the placement within the channel, any water molecule that is within 4.0Å of center of TM core is counted.

## Results

### Homology modeled structures of xCT in two distinct conformations of transition cycle

In the absence of a crystal structure of any cystine transporter, we built homology model of human cystine transporter *xCT* based on crystal structures of transporters that were expected to have similar folds. For building such a model, comparative analysis of xCT sequence using CCTOP server predicted the overall topology of the transporter to consist of 12 transmembrane domains. Based on the region spanning the transmembrane domain region (residues 45 to 512) two templates with 51% sequence similarity were identified for modeling using HHPred: ApcT (broad-specificity proton-coupled amino acid transporter from *M. jannaschii*) and AdiC (an arginine/agmatine antiporters of *E. coli).* This was followed by modeling of the transmembrane (TM) region of xCT using Modeller9v15.

Crystal structures of these two transporters have similar structural core LeuT-fold of APC transporters which consists of two intertwined 5-TM-helix repeats (TMs 1-5 and TMs 6–10) sharing a two-fold inverted pseudo symmetry(95). The two templates are in different conformational state of substrate transport cycle. ApcT structure is reported in substrate free inward facing occluded in pdbid:3GIA. AdiC is reported in substrate free outward facing state in pdbid:5J4I. The overall structure of xCT modeled on the above two templates reveals an overall cylindrical shape with 12 transmembrane helices with short extracellular and cytoplasmic loops and intracellular amino and carboxy termini (Fig. 1). In two models, the 12 TMs are arranged in two intertwined V-shaped inverted repeating units (TMs 1-5 and TMs 6–10), followed by TMs 11 and 12. Our models are in line with the biotinylated experiments that proposed the first topological model for xCT of 12 transmembrane domains with the N and C termini located inside the cell(40). For the inward occluded model, we observe that the intracellular loop IL23 between TM helices 2 and 3 occludes the intracellular site, and this loop is believed to be involved in substrate permeation pathway(40). The modelled structures were then embedded in heterogeneous lipid bilayer and further refined using unrestrained molecular dynamics simulations. Superposition of the simulated model structures with the templates have been shown in Fig SI3. All the models reveal a cavity lined by residues of TM1, TM3, TM6, TM8, and TM10 helices. These five transmembrane helices form the substrate translocation channel and the cavity was assessed to be large enough to accommodate the substrates, and therefore, the probable ligand binding sites (Fig. SI4). Based on the template used and cavities within channel visualizations using HOLE suite of programs, we termed as Model-Cioc and Model-Cout. The former is modeled using ApcT structure (PDB:3GIA), and the latter is modeled using AdiC structure (PDB:5J4I) as templates.

**Fig. 1:**
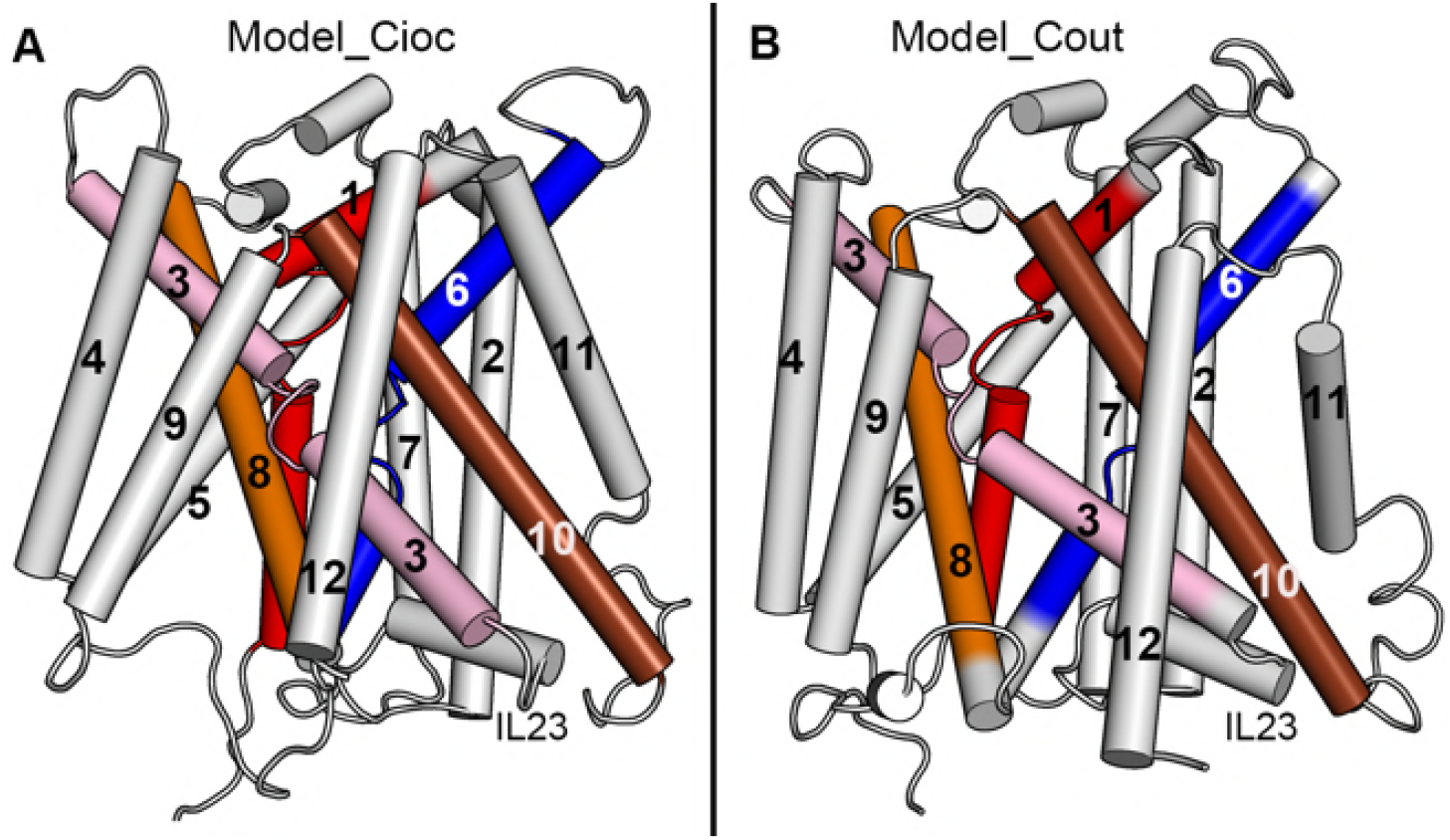
Homology structure of xCT in A. inward facing occluded state Model-Cioc; and B. outward facing state: Model-Cout. The transmembrane helices are marked and the five helices forming probable translocation channel are shown colored. The intracellular loop IL23 is also shown.

### Transition between two modeled xCT conformations using targeted molecular dynamics

xCT is an antiporter and like any other transporters, will have to go through various conformational cycles transporting substrates in and out. We generated two models of xCT transporter in two distinct conformations of substrate transport cycle. The template crystal structures were obtained in absence of ligands, and thus, the models obtained are representative of substrate-free states.

We employed several targeted molecular dynamics (TMD) simulations to drive one conformation to another and the variation of root mean square deviation with respect to time is shown in Fig. SI5. We investigated relative motions of transmembrane domain during TMD simulations using principal component analysis (PCA)(96). We pooled in all the conformations obtained during TMD simulations; and obtained the corresponding eigenmodes and eigenvalues. The eigenmodes associated with the largest eigenvalues have the largest contribution to the dynamics. The first three eigenmodes reflected majority of the motions during the trajectory, accounting for ~75% of the significant fluctuations. We analyzed the time behavior and distribution of the motions performed by the first top modes by projecting the conformations obtained during two cycles of transition: Cycle I and Cycle II. As can be seen from the two-dimensional scatterplots between principal components: PC1, PC2, and PC3 (Fig. 2A); the conformations from two different cycles occupy mirrored conformational subspace as expected from their transitions cycle. To further visualize the significant conformational motions during transition, we generated an interpolated trajectory between the extreme conformations sampled along the first eigenvector or PC1. As can be seen from the porcupine plots along PC1 (Fig. SI6), major changes are observed for the helices TM11 and TM12 along with the intracellular and extracellular loops. The inside helical core forming the probable ligand binding channel showed lesser fluctuations (Fig. 2B), suggesting the role of outer helices in effecting the conformational transition, and probable role of inner helical core in transporter functioning.

**Fig. 2.**
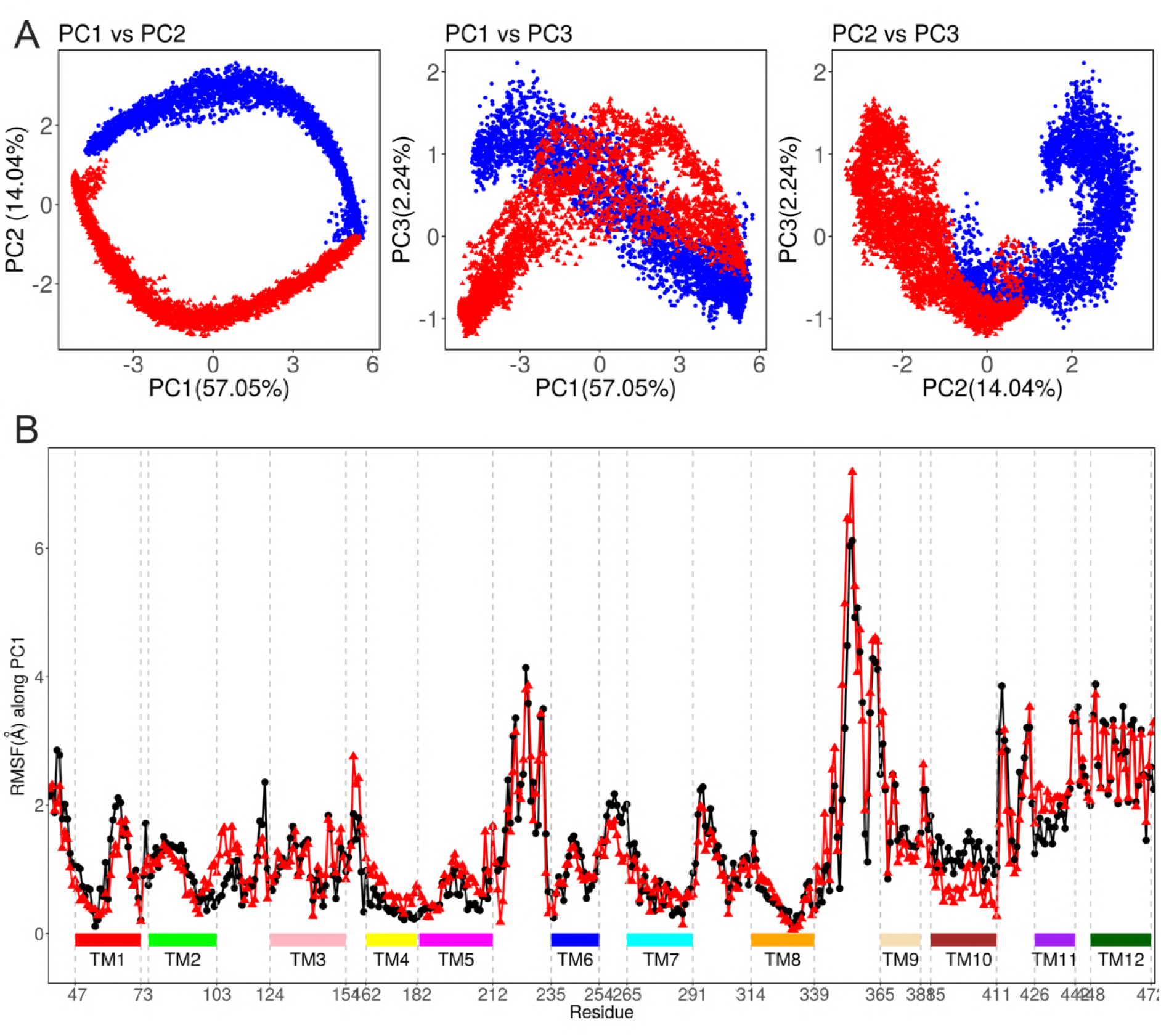
Principal Component Analysis. A. 2D scatterplots of first three eigenvectors as the trajectories are projected onto combined trajectory of TMD simulations runs. Blue color dots correspond to TMD cycle I as transition from Model-Cioc to Model-Cout, and red color triangles corresponds to TMD cycle II as transition from Model-Cout to Model-Cioc. B. Root mean square fluctuations observed for C atom of each residue along the first eigenvector (PC1). Black dots correspond to cycle I and red triangles corresponds to cycle II.

### Probable location of substrate translocation channel in xCT

To elucidate further the role of probable substrate channel in substrate binding and translocation, we carried out simulations in the presence of substrates. We had two options to consider for elucidation of substrate translocation pathway: (a) one to dock the ligands to these modelled structures and then employ end point targeted molecular dynamics; and (b) second to employ the transitional conformations going from one substrate-free conformation to another and then build up the substrate transition pathway by docking the representative structures obtained during the transition with ligands. Owing to the fact that the initial models are based on substrate free conformations lacking the information on substrate binding mode; and the unknown computational time required by substrate to sample the translocation channel during the transition; we opted for second method. We first sampled the transition between substrate free modeled conformations through series of short and long targeted molecular dynamics simulations; and clustered all the conformations into representative structures. We designated these structures as intermediate structures as transition cycle as xCT undergoes transition from inward occluded state to outward state. We docked these intermediate structures with ligands: anionic cystine and glutamate, to view the interactions of ligands during this transition cycle.

### Docking of ligands within the translocation channel of intermediates

We selected ten representative structures from the ensemble of intermediates conformations obtained for both transition cycles and docked with the substrates. Since xCT is an antiporter that exports glutamate and imports cystine, we considered following probable scenarios for substrate translocation from intracellular to extracellular side or *vice versa*:

a. Bimodal transport: xCT undergoes transition from inward-facing to outward facing conformation and transports intracellular glutamate out. This is followed by another transition from outward facing to inward-facing conformation as extracellular cystine is transferred inside the cell.
b. Unimodal transport: xCT transports both cystine and glutamate substrates in opposite directions in one cycle while undergoing transition from inward-facing to outward-facing or outward-facing to inward-facing.

For the first scenario, we docked the intermediates with single substrate either cystine or glutamate and visualized the ligand binding pockets and identified the interacting residues within the transporters. For the latter scenario, we docked these single-substrate bound structures with another substrate and visualized the dual ligand binding interactions.

### Probable bimodal transport of ligands

We analyzed the cavities within intermediate structures using HOLE program and docked anionic cystine or glutamate substrates to these cavities. 500 docked conformations were generated with 50 docked poses for each of 10 intermediate structures, and unique poses were filtered based on autodock vina score and manual inspection. For anionic cystine and glutamate, 45 and 52 unique docked conformations were identified, respectively. These ligand docked structures were then simulated in lipid environment; and the interactions of the ligands with the amino acids were studied. Visualizing all the docked transporter conformations, we observed that the translocation channel for xCT transporter is composed of five transmembrane helices: TM1, TM3, TM6, TM8, and TM10; and ligands were docked within this channel. Based on structural analysis of models and positioning of proteins in membrane (PPM server), the length of translocation channel is ~16.0Å. We further analyzed the placement of ligands within the translocation channel by calculating the z-component of distance between center of mass of ligand and center of mass of translocation channel. Positive value of z-component indicates placement towards the extracellular site and negative value indicate placement towards the intracellular site. We observed the variations of this z-component with respect to the normalized rmsd difference of the translocation channel from the initial modelled structures. The latter will hint whether the transporter structure is close to inward-facing occluded model (Model-Cioc) or outward-facing state (Model-Cout). All the docked conformations were clustered based on these two parameters using density based dbscan clustering method, and obtained representative structures corresponding to unique clusters for ligand bound structures (Fig. 4). The conformations on the far left side of x-axis with negative values closer to −1 are similar to xCT structure modeled in inward-facing occluded state; and the conformations on the far right side of x-axis with positive values closer to +1 are similar to xCT structure modeled in outward facing state. From docking results, we observed that there are certain conformations where ligands bind near the extracellular facing site. No conformations however, were observed where ligands bound near intracellular site. But that may be because one of the end point conformation of targeted molecular dynamics here is in inward facing occluded state, thus preventing the ligands to access further towards the intracellular site.

**Fig. 3.**
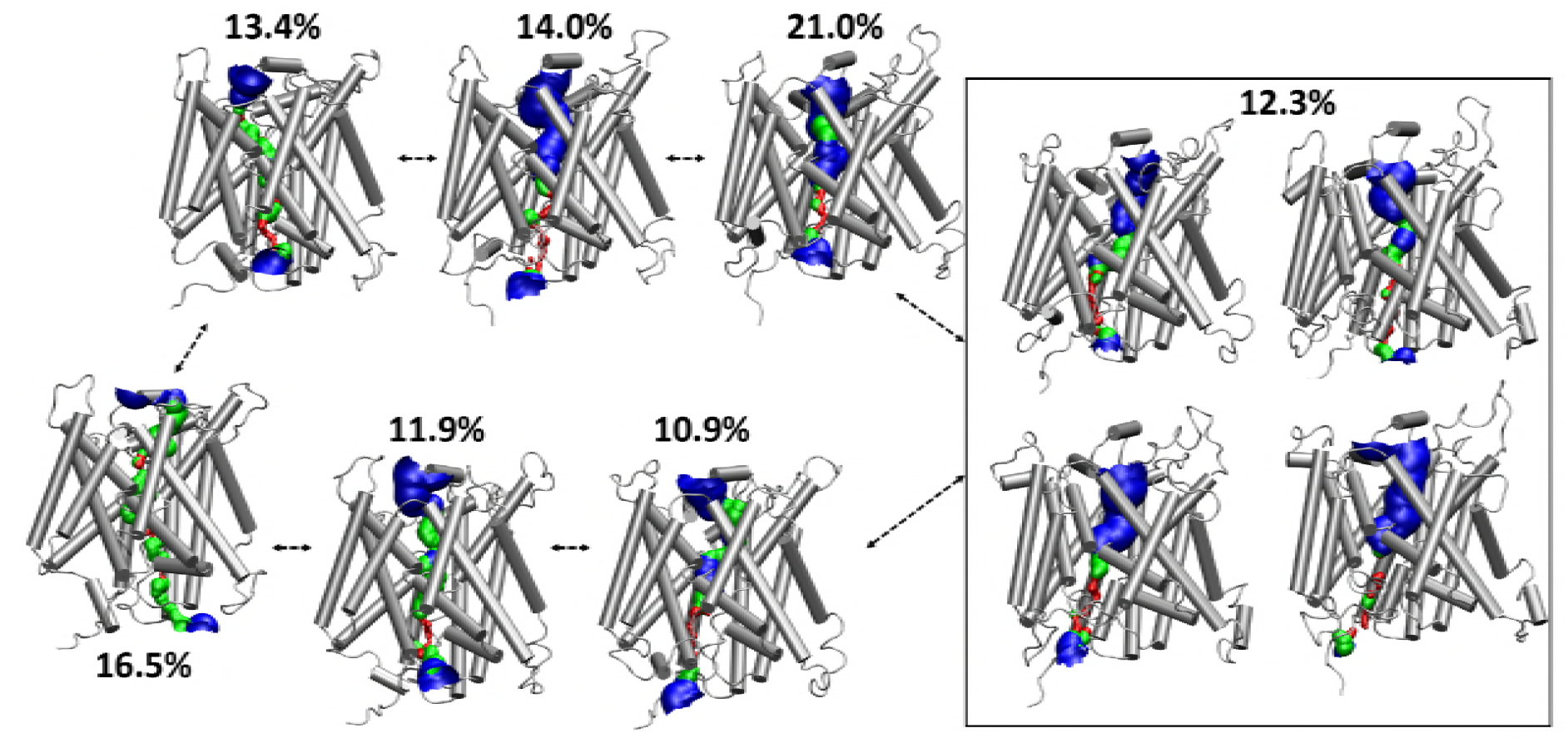
Representative structures as obtained from kmeans clustering of all the conformations obtained during all TMD simulations. The values in percentages show the fraction of conformations forming the cluster corresponding to the respective structures.

**Fig. 4.**
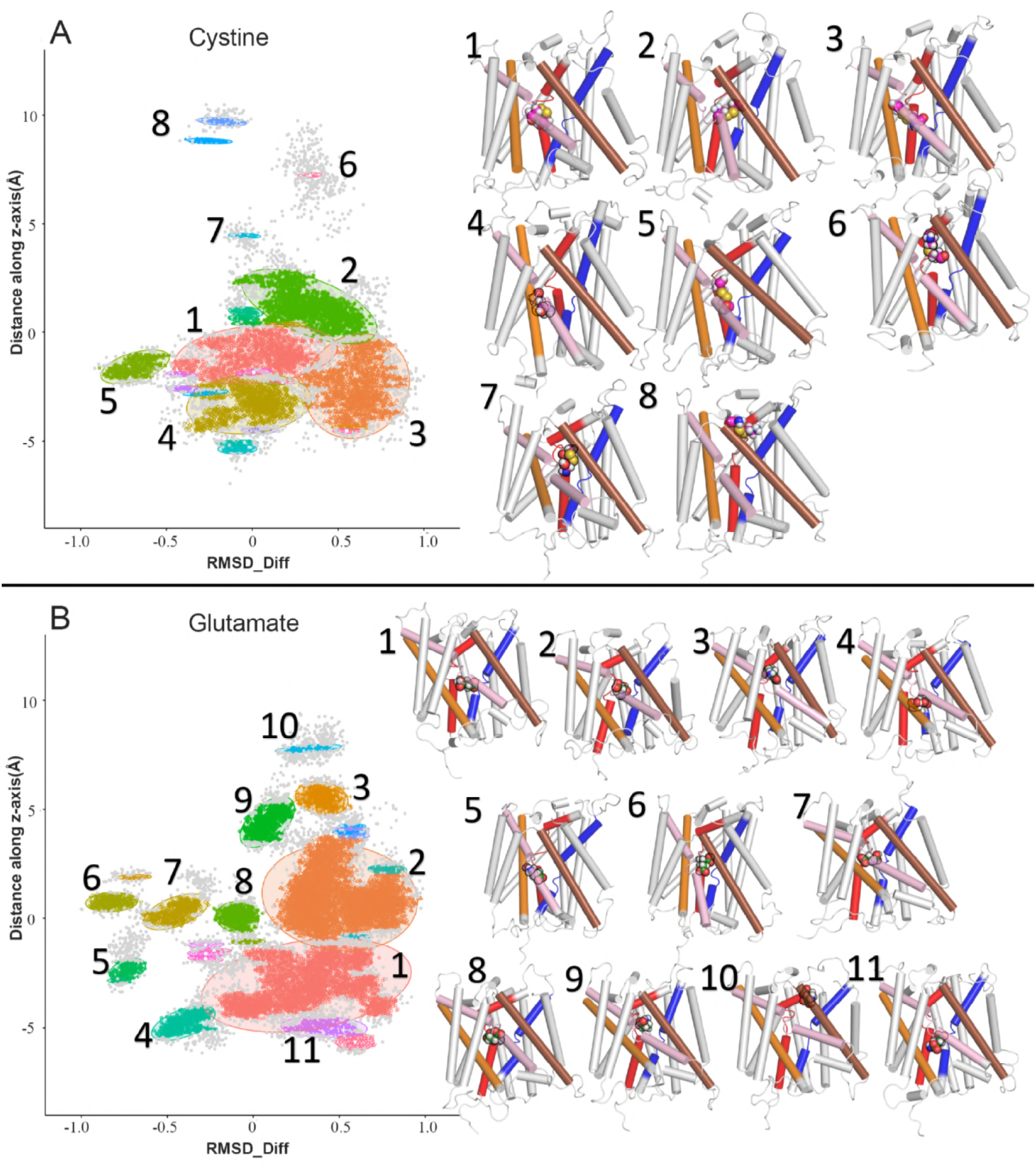
Scatter plots depicting clustering of ligand docked conformations. A. Clustering of conformations docked with anionic cystine, with optimal eps value of 0.075 and minimum points of 50. B. Clustering of conformations docked with glutamate with eps value of 0.07 and minimum points of 50. xCT structures best fit to the centroids of clusters are shown alongside. The helices are colored as in Fig.1, and the ligands are shown in spheres. The outliers are shown as grey spheres.

### Probable unimodal transport of ligands

In a unimodal scenario, extracellular cystine intake is mediated by simultaneous exchange of intracellular glutamate. From the dual docked simulations, we observed that the cavity within the translocation channel is wide enough to bind both the ligands and transport side by side. We clustered the bound conformations based on the z-component of distance between the ligands, and normalized rmsd difference (Fig. 5). The former will indicate the relative position of ligands within the channel, with negative values indicating that glutamate is placed towards the extracellular site with respect to cystine, and positive values indicating that glutamate is placed towards the intracellular site with respect to cystine. We did comparative analysis of conformations of substrate translocation channel sampled with both cystine and glutamate ligands docked with respect to the end point modeled conformations and those observed with either cystine or glutamate docked. The right shift in the conformations along the x-axis (normalized rmsd difference) as observed in Fig 5. suggested that inward facing occluded conformations do not have enough space to bind both ligands. Rest of conformations have enough space to accommodate both ligands. In order to delineate further the bimodal transport, we looked at the interpolated trajectory between two conformations at the extreme ends of cluster analysis. The first conformation is representative of right bottom of cluster (Cluster#9), where cystine is closer to intracellular site and glutamate near to extracellular site. This conformation can be the probable conformation where cystine is transported to intracellular site as glutamate is transported out to extracellular site. The other extreme conformation is representative of upper left cluster (Cluster #8), where glutamate is closer to extracellular site and cystine near to intracellular site. This conformation can be the probable conformation where glutamate is effluxed out while cystine is influxed in. Animation movie shown in movie SM1 indicate the probable simultaneous transport of cystine and glutamate.

**Fig. 5.**
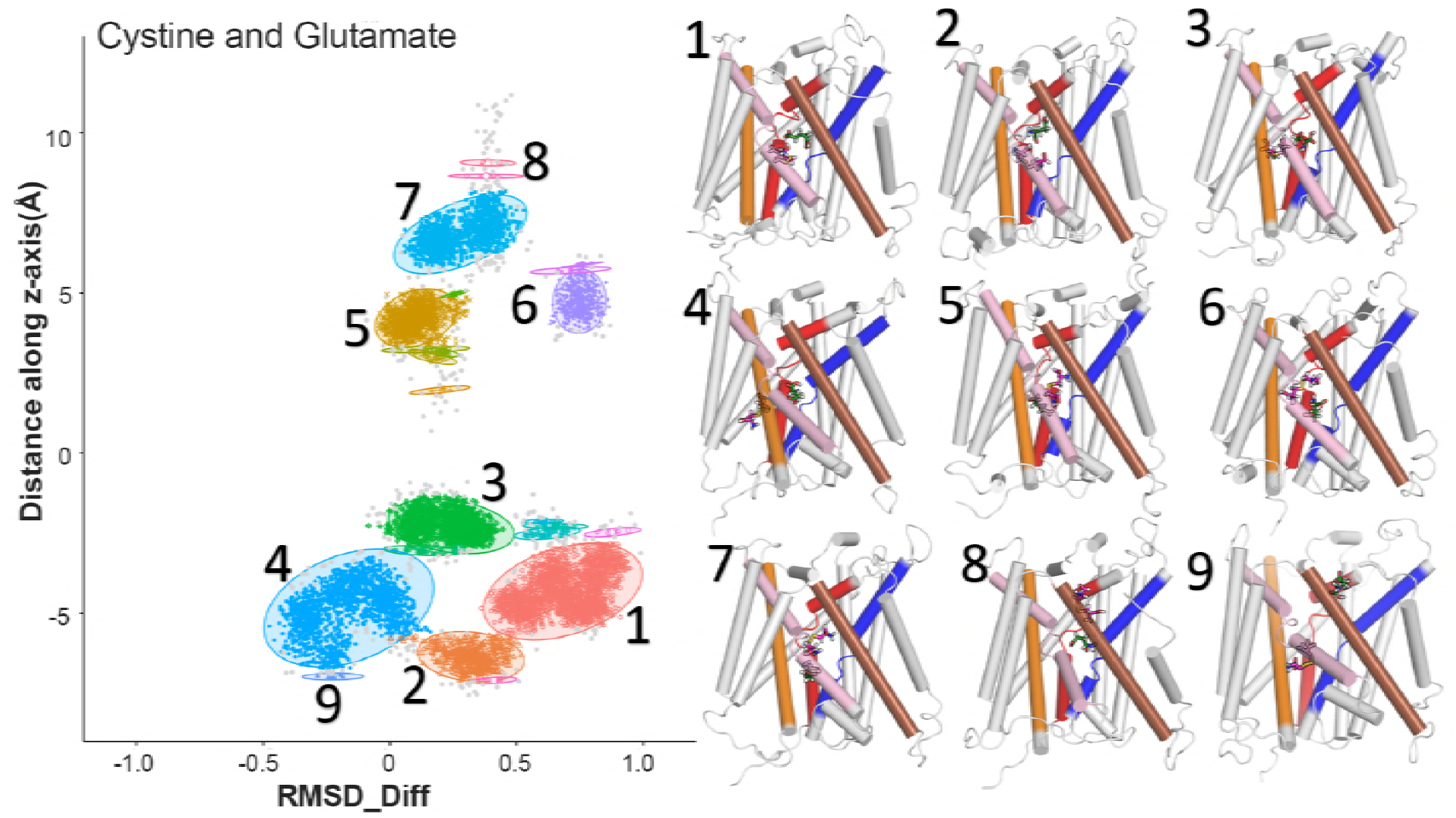
Scatter plots depicting clustering of xCT conformations docked with both cystine and glutamate; with optimal eps values of 0.65 and minimum points of 50. xCT structures best fit to the centroids of clusters are shown alongside. The helices are colored as in Fig. 1, and the ligands are shown in spheres. The outliers are shown as grey spheres.

### Probable amino acids critical for ligand binding

We observed the interactions of the ligands with the amino acids of transporter protein and found that similar residues are mostly involved in interacting both anionic cystine and glutamate (Fig. 6). Most of the interacting residues belong to transmembrane helices: TM1, TM3, TM6, TM8, and TM10. For the intracellular loop (IL2), H110 was proposed to lie close to the permeation pathway, and we do observe that in docked conformations, ligands especially cystine forms weaker interaction with H110. For the extracellular loop (EL1), we observe that two residues Q71 and N72 forms transient interactions with both the ligands, and cystine forms further weak interactions with EL4 with residues L298 to N301. These may act as meeting points for ligands from the extracellular site. Experimental evidence(41) shows that C327 lies close to substrate permeation pathway, and our results do show intermittent interactions of ligands with C327. Experimental mutagenesis data for human xCT is lacking. There is experimental mutagenesis evidence of cystine binding in the homologous fungal cystine transporter, CgCYN1(43); where mutations critical for cystine binding in CgCYN1 were observed; and mapped onto the modelled structure of human xCT. The study suggests that residues F146, A247, G333, L389, and V404 are essential for substrate binding. Our interaction analysis indicated that these residues or their neighboring residues form strong and pertinent interactions with both cystine and glutamate. In addition, we observed that ligands form strong interactions with “G^59^A^60^G^61^” motif present in TM1; and mutations in similar GXG motifs in other transporters led to severe defects(97, 98). Another conserved motif in TMD6: “(F/Y)(S/A/T)(F/Y)xGxx” have been identified critical for transporters within APC family(98). For human xCT, this motif is Y^244^AYAGWF^250^ and is observed to form very strong interactions with both the ligands.

**Fig. 6.**
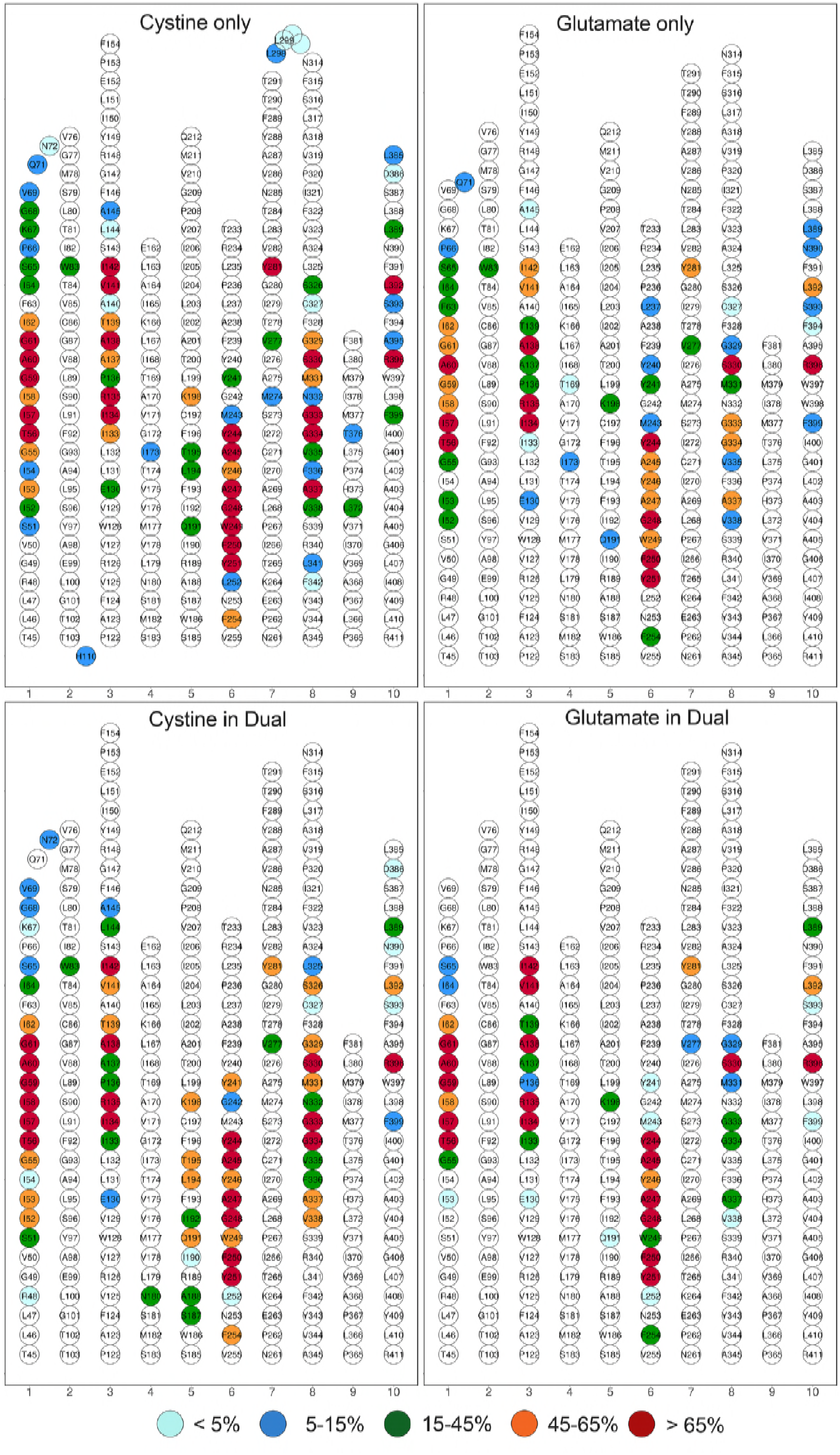
Interactions of ligands with the transmembrane helices. TM helices 11 and 12 are omitted as they do not interact with ligands. Residues that do not interact at all with the ligands are in white circles, and other residues are in colored circles depending on the frequency of their interaction with the ligand. The legend indicates the color corresponding to the frequency of occurrence of interaction observed for all the docked conformations.

### Water molecules within the channel

We observed that during the ligand-docked simulations, the water molecules enter the channel and occupy places along with the ligands. We thus analyzed the total number of molecules within the substrate permeation channel. To ensure the placement within the channel, any water molecule that is within 4.0Å of center of TM core is counted. Figure 7 shows the variation in total number of water molecules within the channel with respect to the normalized rmsd difference. It can be observed that more or less similar number of water molecules are present when only cystine or only glutamate are bound. Conformations towards the inward facing occluded side has a smaller number of water molecules, as less space is available within the channel. Also, when two ligands are bound, relatively less number of water molecules are observed as expected.

**Fig. 7.**
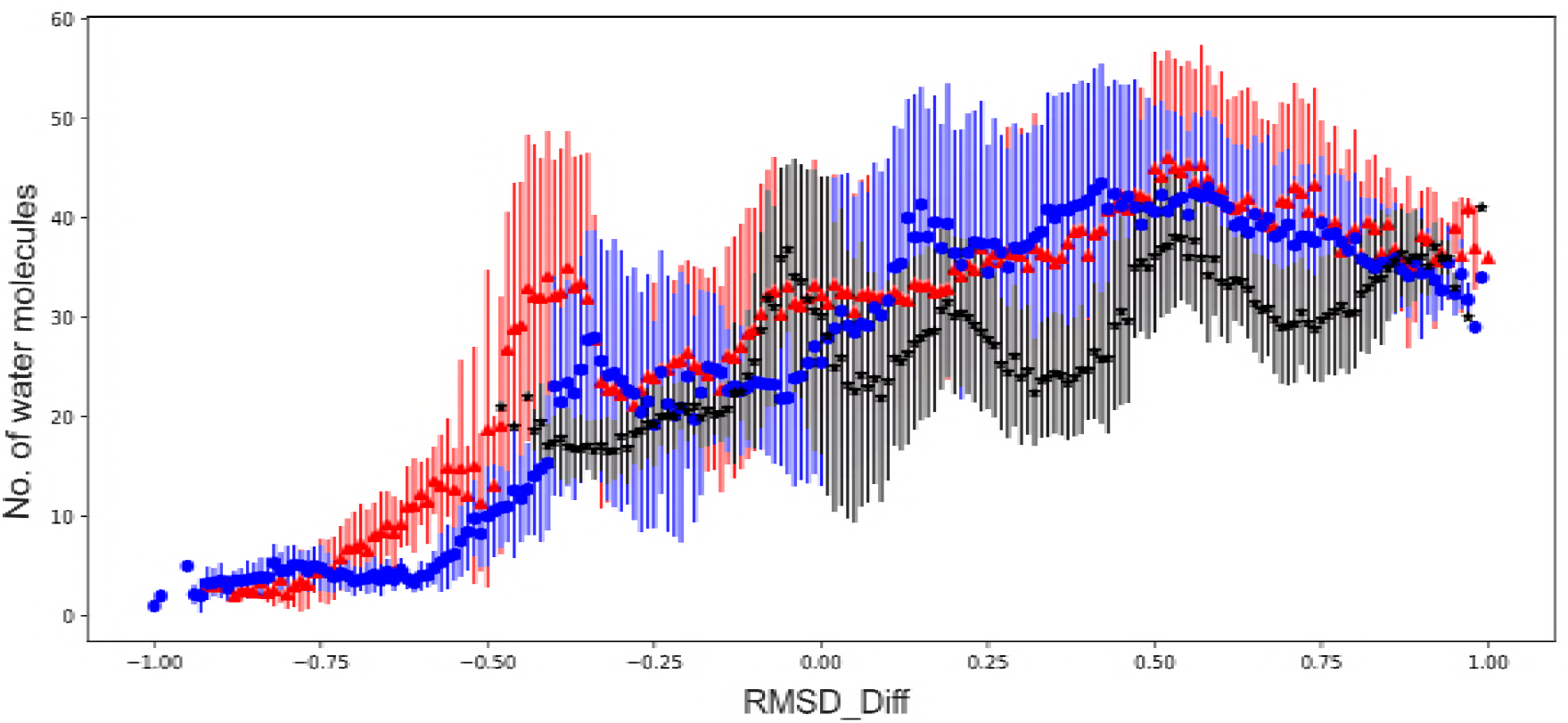
Number of water molecules observed within the translocation channel of ligand docked xCT conformations. Blue colored circles denote water molecules for glutamate-only docked xCT conformations, red colored triangles denote water molecules for cystine-only docked xCT conformations, and black stars denote water molecules for both glutamate and cystine docked xCT conformations.

## Discussion

A transporter undergoes various conformational transitions switching accessibility from extracellular site to intracellular site, and simultaneously transporting the substrates. Human xCT is an antiporter that exchanges intracellular glutamate to take up extracellular cystine. Based on its sequence similarity with other transporters, we generated two human xCT models in two different conformations: inward facing occluded state; and outward facing state. For Model-Cioc, the inward face is observed to be occluded by intracellular loop between helices TM2 and TM3. Both models adopted similar LeuT-fold with 12 transmembrane helices arranged in two intertwined V-shaped inverted repeating units (TMs 1-5 and TMs 6–10), followed by TMs 11 and 12. Comparative analysis of these models suggested presence of substrate translocation channel of cylindrical shape formed by five helices TM1, TM3, TM6, TM8, and TM10.

We further carried out conformational transitions using targeted molecular dynamics simulations from one model to another as end points, and *vice versa.* Series of TMD simulations were run for shorter and longer timescales; and suggested that the majority of the conformational change is effected by the residues in loops and helices TM11 and TM12. Rest of the helices show considerably less effect, indicating the binding cavity for the channel is maintained during the conformational transition. We then identified the intermediates during the transition simulations based on clustering analysis; and docked them with the ligands to understand the ligand binding. For xCT being an antiporter, two scenarios seemed feasible. In first scenario, intracellular glutamate is effluxed out, and then extracellular cystine is taken in; or *vice versa.* In second scenario, as intracellular glutamate in effluxed out, extracellular cystine is simultaneously taken in. So, we did two docking studies: first with single ligands, and second with both the ligands.

All the docked conformations were simulated to allow ligands to explore conformational space within the channel. These conformations were then clustered to identify the conformations that uniquely represent the placement of ligand within the channel as the xCT adopts certain conformation. There were conformations observed where the ligands are placed near the extracellular site. In case of single ligand binding (bimodal transport), these conformations can be marked as; when the extracellular cystine is taken in (as represented by Cluster#6 in Fig. 4A) and when the intracellular glutamate is transported out (as represented by Cluster#8 in Fig. 4B). Since xCT is modeled in intracellular occluded state, we observe that cystine and glutamate interacts with IL23 on their way from extracellular and intracellular sites as represented by Cluster#5 in Figs. 4A and 4B, respectively. In case of dual ligand binding (unimodal transport), it seems that the inward occluded model of xCT does not have enough space to bind both ligands and suggested slight opening up of channel when both ligands are present. We however, do see feasible transition as glutamate and cystine are transported out and in, simultaneously. However, we cannot rule out the fact that transition from inward facing occluded state to inward facing open conformation may facilitate further the simultaneous transport of anionic cystine and glutamate across the membrane. This will require more investigation of transition dynamics going from occluded to open state and is a subject of further studies beyond this work.

We also investigated the interactions of ligand with the xCT. Both ligands seem to bind similar residues, specifically those belonging to helices TM1, TM3, TM6, TM8, and TM10. These interactions were then categorized based on their occurrence during all the ligand bound simulations, and the critical residues for ligand binding were observed. Though, human xCT mutagenesis data is lacking, but mutagenesis study of homologous fungal transporter, CgCYN(43) suggested that mutating similar residues in CgCYN resulted in severe defects in transporter functioning. The proximity of C327 belonging to TM8 to the substrate permeation pathway was observed experimentally(41), and we do observe its intermittent interactions with the ligands in our docked conformations. We also observed the presence of water molecules within the channel along with the ligands.

Combining the transition dynamics and docking studies, we have investigated the ligand translocation mechanism for human cystine transporter, xCT as it undergoes conformational transition from extracellular open to intracellular occluded state. Our studies indicate that there is feasibility of both the scenarios when the ligands can be transported in unimodal mode as well as bimodal mode. To explore the complete scenario, more experimental structures or data are required to investigate further about the intracellular open state of xCT.

## Conclusion

We have investigated the transition of xCT between two conformations: inward occluded state and outward open state. The conformational changes during the dynamics were studied, and intermediate structures were identified. The latter were used for docking studies where anionic cystine and glutamate were docked either alone or in combination. We observed that the ligands bind within the translocation channel formed by helices TM1, 3, 6, 8 and 10; and similar residues interact with the ligands. These residues map well to the experimentally known important residues important for ligand binding in human xCT or homologous fungal cystine transporter, CgCYN.

## Acknowledgements

MS thanks Department of Science and Technology (DST), India for INSPIRE Award and research grant (IFA14-CH-165).

## Supplementary Information

Supplementary document consists of figures SI1-SI6.

## Contributions

M.S. conceived, designed, and performed the experiments. A.C.R performed initial experiments. M.S. analyzed the data and wrote the paper. M.S. and A.C.R contributed the literature materials and reviewed the manuscript.

